# Season-specific dominance broadly stabilizes polymorphism under symmetric and asymmetric multivoltinism

**DOI:** 10.1101/2023.11.20.567918

**Authors:** Evgeny Brud

**Affiliations:** Department of Biological Sciences, North Carolina State University, Raleigh, NC, USA; Department of Biology, University of Pennsylvania, Philadelphia, PA, USA

## Abstract

Seasonality causes intraannual fitness changes in multivoltine populations (defined as having multiple generations per year). While it is well-known that seasonally balanced polymorphism is established by overdominance in geometric mean fitness, an unsettled aspect of the deterministic theory is the relative contribution of various season-specific dominance mechanisms to the potential for polymorphism. In particular, the relative importance of seasonally-reversing and non-reversing schemes remains unclear. Here I analyze the parameter space for the discrete generation two-season multivoltine model and conclude that, in general, a substantial fraction of stabilizing schemes are non-reversing with the season (∼25-50%). In addition, I derive the approximate equilibrium allele frequency cycle under bivoltinism, and find that the amplitude of allelic oscillation is maximized by non-reversing dominance if the selection coefficients are roughly symmetric. Lastly, I derive conditions for the intralocus evolution of dominance. These predict a long-term trend toward maximally beneficial reversal. Overall, the results counter the disproportionate emphasis placed on dominance reversal as a stabilizing mechanism and clarify that non-reversing dominance is expected to frequently characterize seasonally fluctuating alleles under both weak and strong selection, especially in their early history. I conclude that seasonally alternating selection regimes are easily able to maintain allelic variation without restrictive assumptions on either selection coefficients or dominance parameters.

## Introduction

The discovery of allelic oscillations in temperate *Drosophila melanogaster*, where hundreds of biallelic sites spread across the genome display seasonal changes in allele frequency of ∼4-20% (Bergland et al. 2014, Machado et al. 2021) has renewed interest in cyclic regimes of natural selection. The simplest model of cyclical selection is the deterministic bivoltine model with non-overlapping generations and a two-season selection regime. Levins (1968) remarked on the perpetual mismatch of selected-traits and seasonal environments in such populations: the winter generation is the offspring of parents selected for traits favorable in the summer, and conversely. Abrupt seasonal switches in populations with >2 generations per year also incur a deleterious load, but at least this situation permits within-season adaptation for those generations experiencing constant selection. Levels of genetic variation in either kind of multivoltine population (defined as >1 gens/year) must partly depend on the prevalence of alleles whose effect is to increase fitness in one season while decreasing fitness in the other (seasonally antagonistic pleiotropy); this owes to the subset of such antagonisms that express overdominance with respect to geometric mean fitness. Classic theory on fluctuating selection in diploids established this criterion (geometric mean overdominance) as a sufficient condition for protected polymorphism (Haldane and Jayakar 1963, Hoekstra 1975). (While complete dominance can also establish variation, the conditions on selection parameters are restrictive). Since cycles of fitness changes are a special class of fluctuating selection (a broader term that includes models with randomly-varying fitnesses), the criterion of geometric mean overdominance carries over as a sufficient condition for stable polymorphism under cyclical selection. This condition conveys that even in the absence of overdominance within a generation, a time-averaged heterozygote advantage can emerge over longer time spans. For multivoltinism, the focal comparison is the array of annual genotypic fitnesses.

More recent work on two-season models (Wittmann et al. 2017, Bertram and Masel 2019) decomposes fitness into season-specific components: selection coefficients (s_1_, s_2_) and dominance coefficients (h_1_, h_2_); in particular, this work investigates the contribution of dominance to the stabilization of polymorphism. Wittmann et al. (2017) conclude from their multilocus modeling that beneficial reversal of dominance, where the seasonally-maladapted allele is always partially or completely recessive, is a powerful stabilizing mechanism for genome-wide polymorphism under either multiplicative non-epistasis or diminishing returns epistasis. In their single-locus treatment, Bertram and Masel (2019) similarly conclude that dominance reversal stabilizes alleles across a broad range of selection intensity; incomplete dominance and additivity preferentially stabilize alleles with large effects on seasonal fitness, and therefore weakly oscillating alleles are expected to be excluded from the genetic architecture of seasonal phenotypes under these special cases, in contrast to dominance reversal.

Previous analyses tended to focus on contrasting special relations for the dominance parameters. Wittmann et al. (2017) assumed symmetric dominance reversal (h_1_ = h_2_) for the analytical results and much of the simulations (except for their figure 10). In their analysis of protected polymorphism, Bertram and Masel (2019, except for their equation 4) compared symmeric dominance reversal to the complement relation (h_1_=1-h_2_, ‘cumulative overdominance,’ Dempster 1955), which includes additivity as a special case. A treatment of generalized two-season dominance ought to clarify the following: (1) Is reversal of dominance ever necessary for the persistence of polymorphism? and (2) Is the exclusion of weakly selected polymorphism typical of broader schemes of incomplete dominance (for which allelic dominance is maintained in direction only, rather than maintained in both direction and magnitude)? These amount to considering the question: *Does season-specific dominance broadly stabilize allelic variation?* In addition, I investigate stability on a longer time scale, and ask: *Subsequent to the establishment of polymorphism, is dominance expected to evolve in a particular direction?*

To attack these questions, I assume a model with a seasonal fitness cycle in which selection occurs for *m* generations of season 1, followed by *n* generations of season 2 (Table 1). I obtain conditions on season-specific dominance for establishing a persistent allelic oscillation, assuming either (a) an equal number generations per season (*m* = *n*) or (b) that the ratio of generation numbers equals *m* : 1 (for *m* > 1). I also obtain (c) the equilibrium allele frequency cycle for *m* = *n* = 1 (bivoltinism), and (d) conditions for the spread of intralocus dominance modifiers into a resident bivoltine equilibrium.

**Table 1.** Fitnesses of diploid genotypes under two-season selection.

|  | AA | Aa | aa |
| --- | --- | --- | --- |
| Season 1<br>($m$ gens) | 1 | $1 - h_1 s_1$ | $1 - s_1$ |
| Season 2<br>($n$ gens) | $1 - s_2$ | $1 - h_2 s_2$ | 1 |

## Model and Results

Consider a multivoltine population with discrete non-overlapping generations subject to the selection regime in Table 1, where the fitness parameters (h_1_, s_1_, h_2_, s_2_) at a biallelic locus are bounded by [0,1]. It is known that a sufficient condition for the establishment of protected polymorphism under temporally varying selection is geometric mean overdominance (Haldane and Jayaker 1963). If W(·) represents the annual fitness, this condition for seasonal fitness cycles is equivalent to:

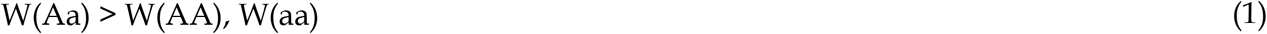

where the rank order of homozygous annual fitnesses is left unspecified. Reformulating Eqn 1 with the season-specific parameters of Table 1 gives:

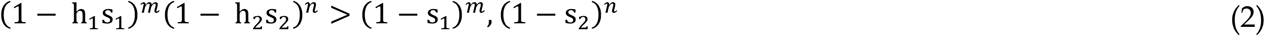

### Symmetric multivoltinism

Suppose that the generations are evenly divided between seasons (*m*=*n*). Then, without loss of generality, we may assume s_1_ ≤ s_2_ and so Eqn 2 simplifies to:

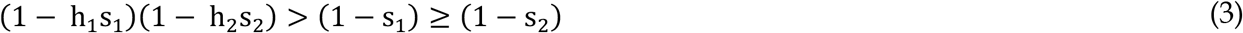

For some (s_1_, s_2_), Eqn 3 reduces to the region *P* on the unit square of dominance parameters:

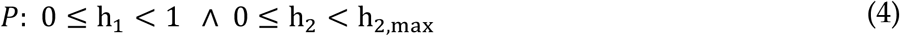

where 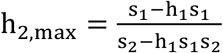.

An equivalent representation of *P* is given by:

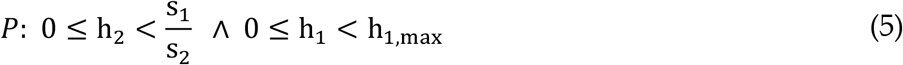

where 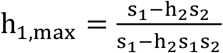. The h_2,max_ and h_1,max_ are concave, monotonically decreasing functions of h_1_ and h_2_, respectively.

*P* includes all (h_1_,h_2_) below (h_1_,h_max,_), or equivalently all (h_1_,h_2_) to the left of (h_1,max_,h_2_) (Figure 1). The area of *P* corresponds to the probability that random sampling from the bivariate uniform distribution bounded by [0,1]^2^ yields a dominance scheme that confers protected polymorphism, i.e. the ‘potential for polymorphism’ (Asmussen and Basnayake 1990, Trotter and Spencer 2007) conditional on biallelism. Integrating h_2,max_ across the range of permissible h_1_ yields the area formula:

**Figure 1.**
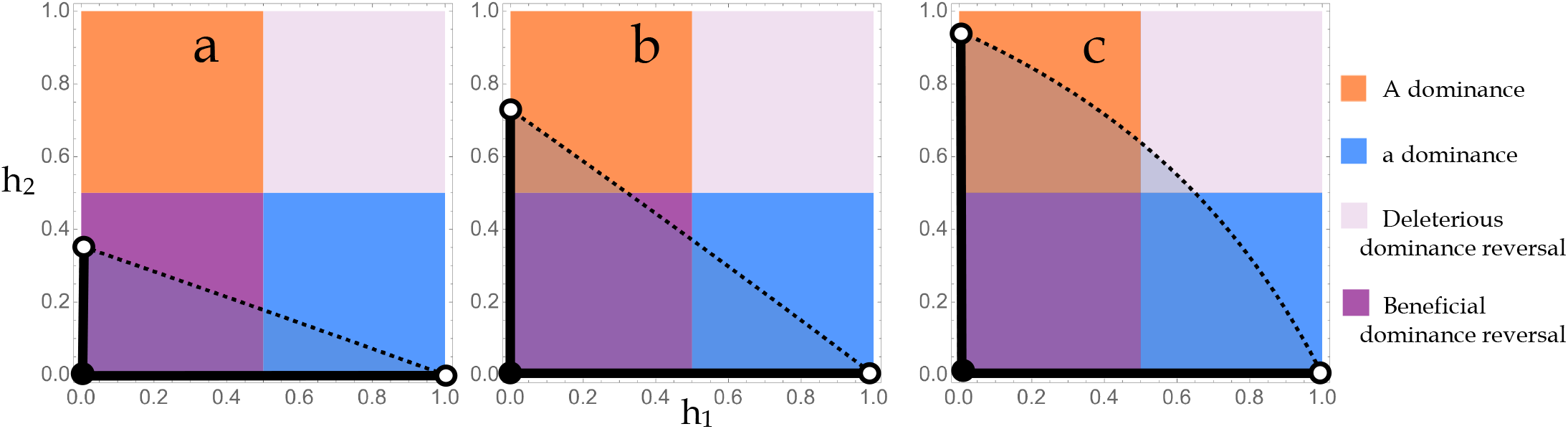
Regions of the dominance unit square. (a-c) The four major patterns of season-specific dominance correspond to a unique quadrant on the unit square of (h_1_, h_2_) values. For arbitrary (s_1_, s_2_), the region *P* conferring protected polymorphism intersects 2, 3, or 4 quadrants as determined by the three cases: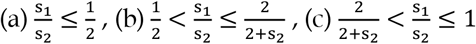 . The h_2_-intercept is the ratio of selection coefficients.

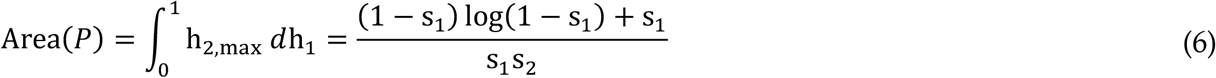

Area(*P*) is considerable (10-50%) for the vast majority of (s_1_,s_2_) combinations (Figure 2a). Increasing asymmetry of the selection coefficients reduces the fraction of dominance schemes in *P* and thereby lowers the potential for polymorphism.

**Figure 2.**
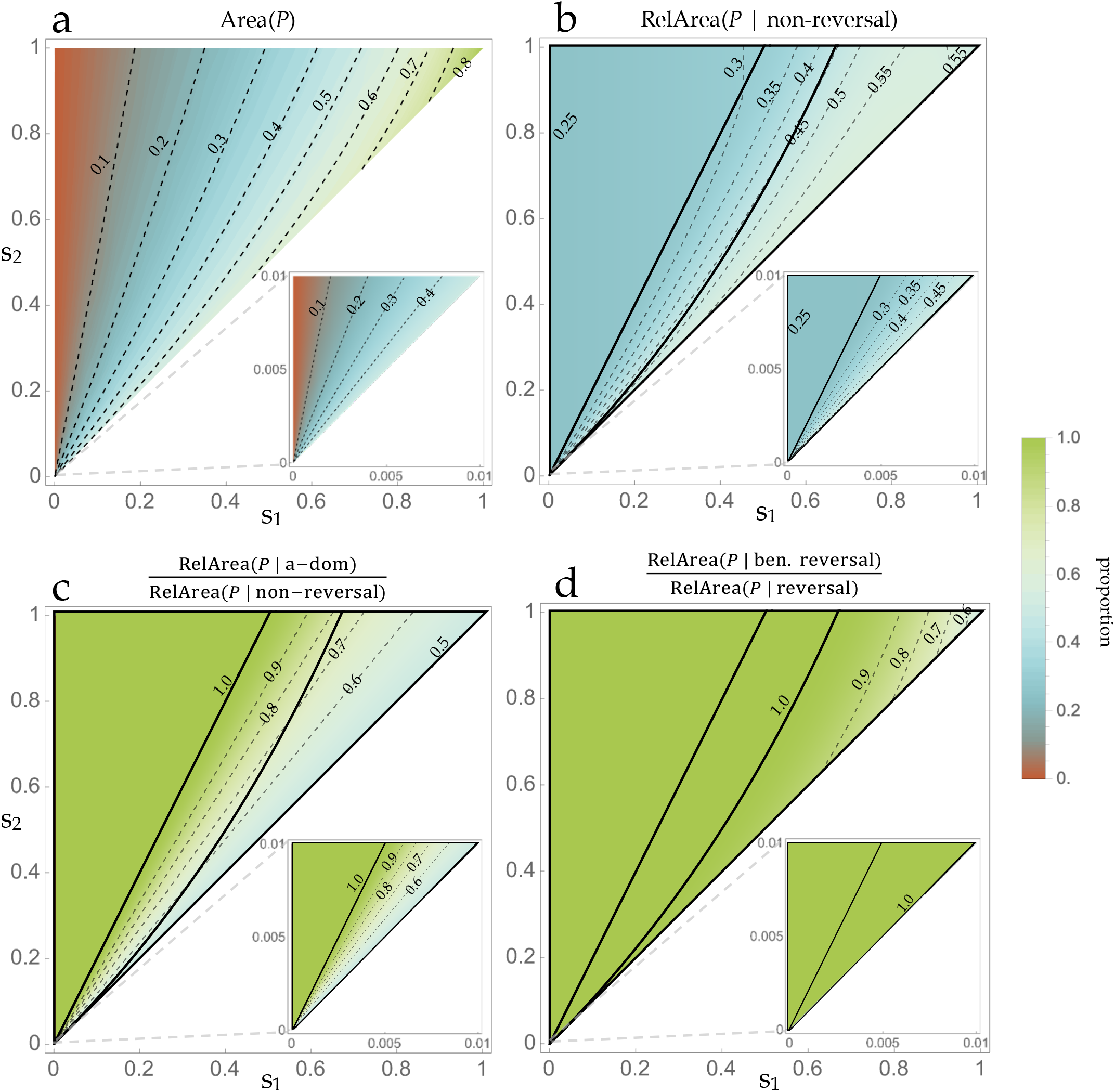
The potential for seasonally balanced polymorphism with equal generations per season (*m* = *n*). (a-d) Contour plots describe the size and quadrant-specific distribution of the region *P*. The blank area owes to the assumption s_1_ ≤ s_2_. (a) Area(*P*), each point corresponds to the fraction of all possible (h_1_,h_2_) schemes that comport with geometric mean overdominance under equal generations per season. For small selection coefficients (inset), the range of areas is restricted to the approximate interval (0, 0.5). (b-d) Bold lines delimit cases i-iii (left to right). For small selection coefficients (inset), case iii is virtually absent. See Box A1 for the definition of the relative area function (RelArea). (b) Fraction of *P* that lies in non-reversing quadrants. (c) Fraction of *P* in the a-dominance quadrant, relative to the fraction of *P* lying in non-reversing quadrants. (d) Fraction of *P* in the beneficial reversal quadrant, relative to the fraction of *P* lying in reversing quadrants.

Area(*P*) sums the contribution over all possible dominance schemes. The relative contribution owing to each major class of season-specific dominance (beneficial reversal, deleterious reversal, generalized A-allele dominance, and generalized a-allele dominance) is determined by the distribution of *P* across the quadrants of the unit square (Appendix I). This distribution is described by three cases, distinguished by whether *P* intersects 2, 3, or 4 quadrants. *Case* i: If the ratio of selection coefficients 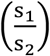 is less than or equal to one-half, *P* is comprised of subregions lying in the quadrants corresponding to beneficial reversal and a-allele dominance. *Case* ii: For 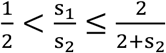, *P* is comprised of subregions in the two non-reversing quadrants and beneficial reversal. *Case* iii: For 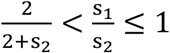, *P* intersects all four quadrants, subsuming the entirety of beneficial reversal.

In general, the relative contribution of non-reversing dominance to the stabilization of polymorphism is substantial (∼25-50%; Figure 2b). With a greater than two-fold asymmetry in selection coefficients (case i), the non-reversing contribution owes entirely to dominance of the a-allele (Figure 2c); this is sensible given that generalized a-allele dominance confers reduced selection against heterozygotes as compared to generalized dominance of the A-allele (Table 1). Greater parity of selection coefficients (cases ii-iii) allows for dominance of the A-allele to contribute to the area of *P* (up to 50% of the non-reversing region, as occurs for s_1_=s_2_), and this applies for general selection intensity (Figure 2c). For the reversing quadrants, it is only under especially strong selection (on the order of 20% or greater viability reduction) that deleterious reversal becomes an important contributor to polymorphism (Figure 2d). Importantly, both a-allele dominance and beneficial reversal are *general stabilizing schemes* in that they both commonly instantiate geometric mean overdominance for selection coefficients of any intensity (so long as the s_1_, s_2_ are not grossly asymmetric). The remaining two schemes (A-allele dominance and deleterious reversal) are specialized, with the former stabilizing polymorphism so long as the ratio of selection coefficients is greater than one-half, and the latter substantively contributing to polymorphic stability only when selection coefficients are near-parity *and* have a strong magnitude. [Recall that s_1_ (the selection coefficient against the a-allele in season 1) was assumed to be less than s_2_ (the selection coefficient against the A-allele in season 2), which explains the differential results between A-and a-allele dominance].

### Asymmetric multivoltinism

Suppose that the generation numbers *m, n* conform to the ratio *m* : 1. Eqn 2 then simplifies to the two distinct possibilities for annual overdominance:

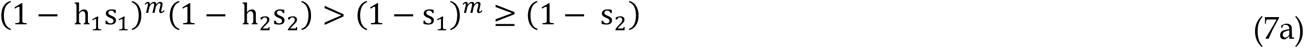

or

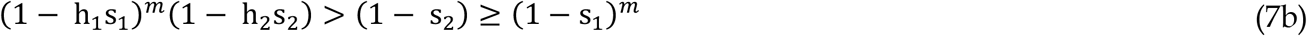

depending on the rank order of homozygous annual fitnesses. (No assumptions are placed on s_1_, s_2_).

Given (s_1_, s_2_,*m*) satisfying the righthand inequality of condition 7a, the region *Q*_*m*_ on the unit square gives the stabilizing dominance parameters:

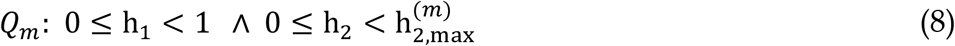

where 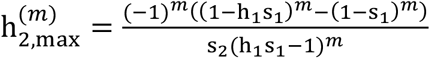.

Integrating 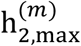 across the range of permissible h_1_ yields the area formula:

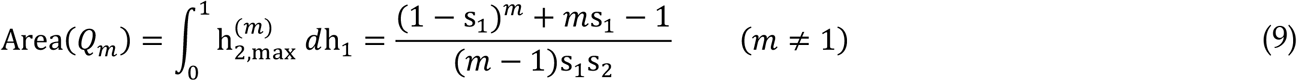

Given (s_1_, s_2_,*m*) satisfying the righthand inequality of condition 7b, the region *R*_*m*_ of stabilizing dominance parameters is equal to:

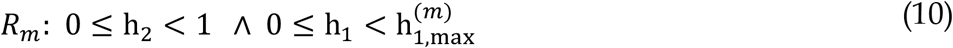

where 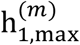 is the smallest real root of the polynomial:

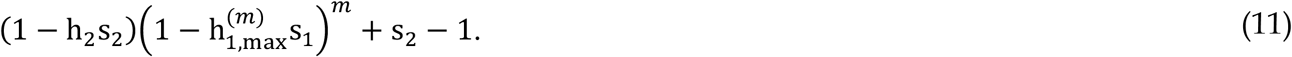

Integrating 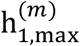 across the range of permissible h_2_ yields the area formula:

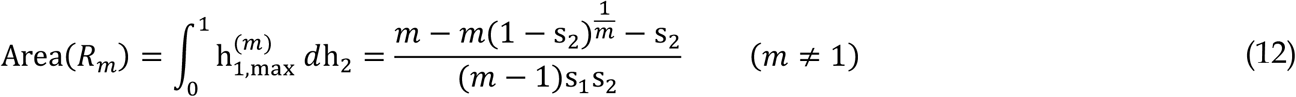

The region *P*_*m*_ determines the overall potential for biallelic polymorphism under an *m*:1 asymmetric distribution of generations between seasons, and is defined piecewise:

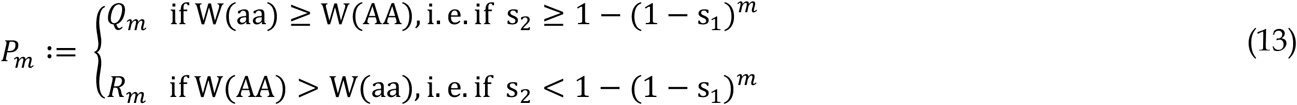

Increasing the asymmetry of multivoltinism (i.e. increasing *m*) constrains the possible (s_1_,s_2_) combinations that are consistent with *Q*_*m*_ and the fitness inequalities in (7a), but simultaneously expands the (s_1_,s_2_) combinations that are consistent with *R*_*m*_ and the fitness inequalities in condition (7b) (Figure 3). Area(*P*_*m*_) is sizeable (∼10-50%) for the great majority of season-specific selection coefficients and so asymmetric generation numbers do not impose severe improbabilities of sampling a stabilizing (h_1_,h_2_) scheme. For small selection coefficients, the results are essentially identical to the Area(*P*) result under symmetric multivoltinism when considering the space of proportionate (s_1_,s_2_) combinations for which 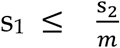 (Figure 3, bottom panels; compare to Figure 2a, inset).

**Figure 3.**
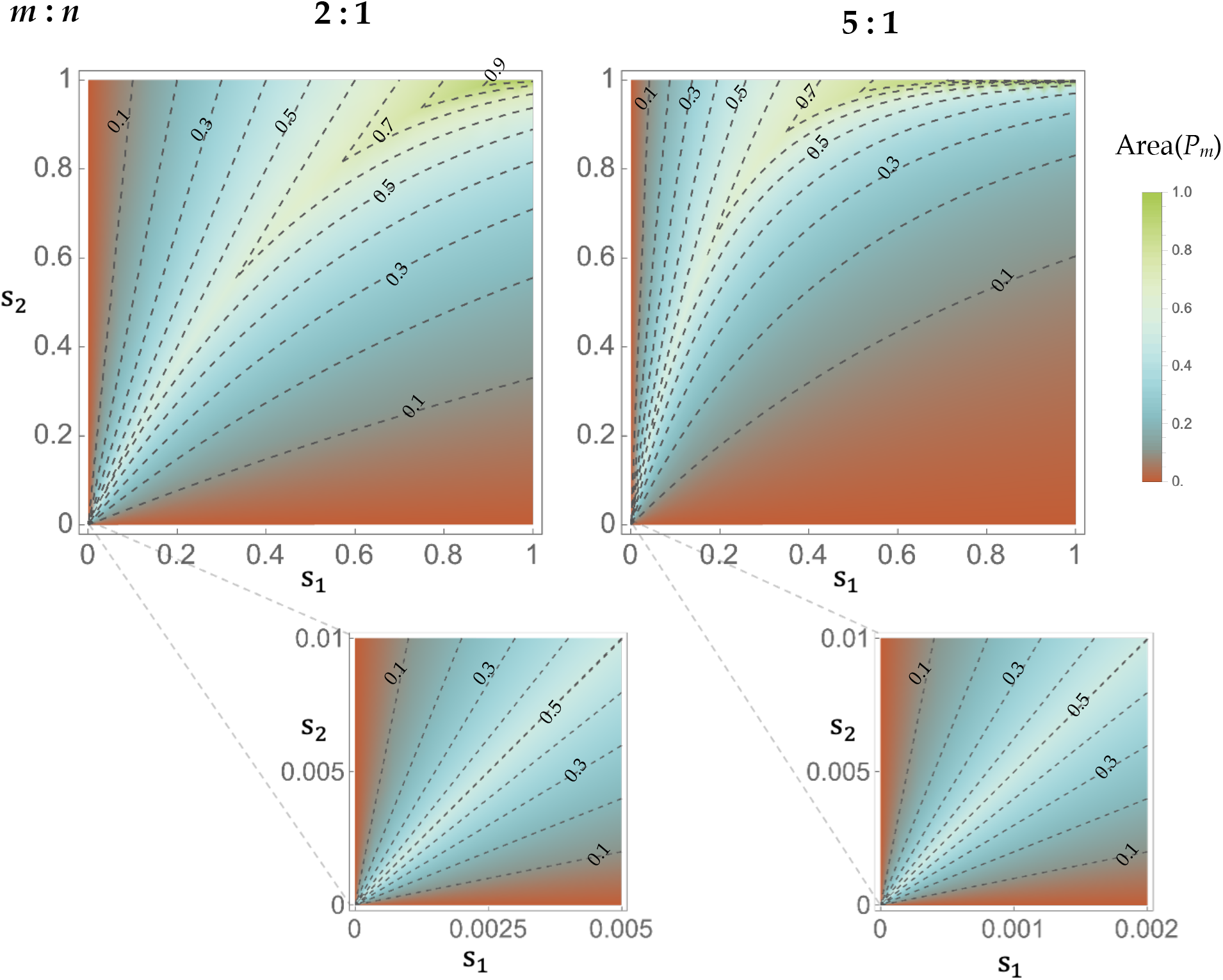
The potential for biallelic polymorphism under an asymmetric ratio of generation numbers. Contour plots give the area of the region *P*_*m*_ assuming the indicated ratios of generations per season. For small selection coefficients (bottom), note that the scale of the axes reflects the generation ratio.

The quadrant-specific contributions can be obtained numerically for any combination of generation numbers. For each case of *m* ∈ (1,2,5,10) and *n* = 1, I obtained a sample of 10^5^ random points from the region described by Eqn 7, assuming three regimes of selection intensity (s_1_,s_2_ ranges: 0-0.1% weak, 0.1-10% moderate, and 10-100% strong), repeated for each rank order of the homozygous annual fitnesses (conditions 7a-b). Each point was then categorized by its dominance quadrant. The resulting distribution of stabilizing dominance schemes is nearly identical between weak and moderate ranges (for those sets sharing *m* values and having the same homozygous fitness ordering) (Figure S1). For W(aa)>W(AA), the weak-to-moderate distribution is notably insensitive to *m*, and is approximately equal to 65%(b): 25%(a): 10%(A): 0%(d) (b: beneficial reversal; a: a-allele dominance; A: A-allele dominance; d: deleterious reversal). For W(AA)>W(aa), the weak-to-moderate distribution is approximately 65-72%(b): 2.5-10%(a): 25.0-25.4%(A): 0%(d). In this latter condition, the contributions of beneficial reversal and a-allele dominance depend on the number of season 1 generations, with increasing *m* modestly expanding the scope of beneficial reversal and concomitantly decreasing that of a-allele dominance. With strong selection coefficients, the distributions are: 41-55%(b): 29.2-29.5%(a): 13-25%(A): 2-4%(d) assuming W(aa)>W(AA), and 55-63%(b): 4-13%(a): 30-32%(A):0.5-2%(d) assuming W(AA)>W(aa). Increasing *m* expands the scope of beneficial reversal under W(aa)>W(AA), but has the opposite effect under W(AA)>W(aa). Under *m=10* and strong selection, the contribution of a-allele dominance shrinks to a mere 4% when the aa homozygote is less fit on an annual basis (Figure S1).

### Allele frequency oscillation under bivoltinism

Consider a population subject to the fitness cycle in Table 1 with *m* = *n* = 1. Assuming discrete non-overlapping generations, the deterministic evolution of allele frequencies in a bivoltine population is described by the equations in Appendix 2.

For small selection coefficients, the relative annual fitness of the a-allele (w_a_) is approximately equal to

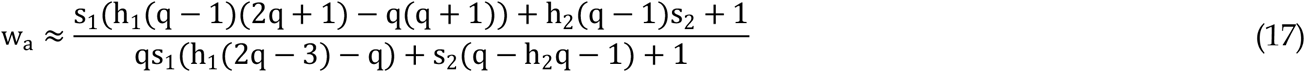

where q is the a-frequency. Solving for w_a_ = 1, I obtain the approximate equilibrium a-frequency at the start of season 1 (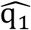, Figure S2a-c):

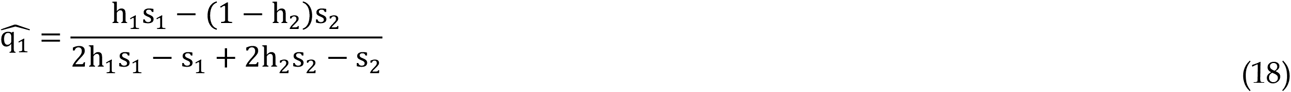

As a matter of fact, 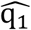 is the classical equilibrium for heterozygote advantage (Fisher 1922) as applied to the annual fitnesses and is equivalent to the expectation under two-trait antagonistic pleiotropy, in which the traits interact additively with respect to viability (Rose 1982). This makes sense since the assumption of small selection coefficients flattens the distinction between arithmetic and geometric mean fitnesses. A unique feature of seasonally balanced selection that transcends constant-selection models of heterozygote advantage is the *allele frequency cycle* implied by the alternation of fitnesses. The approximate magnitude of this oscillation can be derived by first solving for the a-allele frequency at the start of season 2,

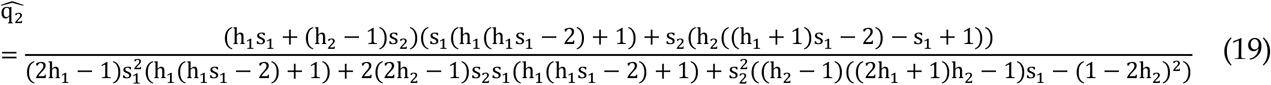

Then the change in allele frequencies between seasons provides the oscillation:

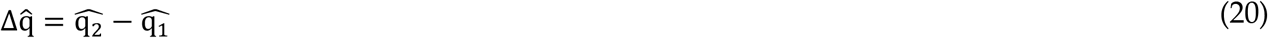

If the selection coefficients are near-parity, 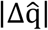 is maximized in the A-dominance quadrant of the unit square (Figure 4a). More disparate selection coefficients result in 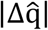 being maximized in the beneficial reversal region (Figure 4c). This pattern also holds under strong selection (Figure 4d-f). Allele frequencies at equilibrium imply considerable levels of heterozygosity unless selection coefficients are especially disparate and/or the dominance scheme lies too close to the maximal boundary of the *P* region (Figure S1).

**Figure 4.**
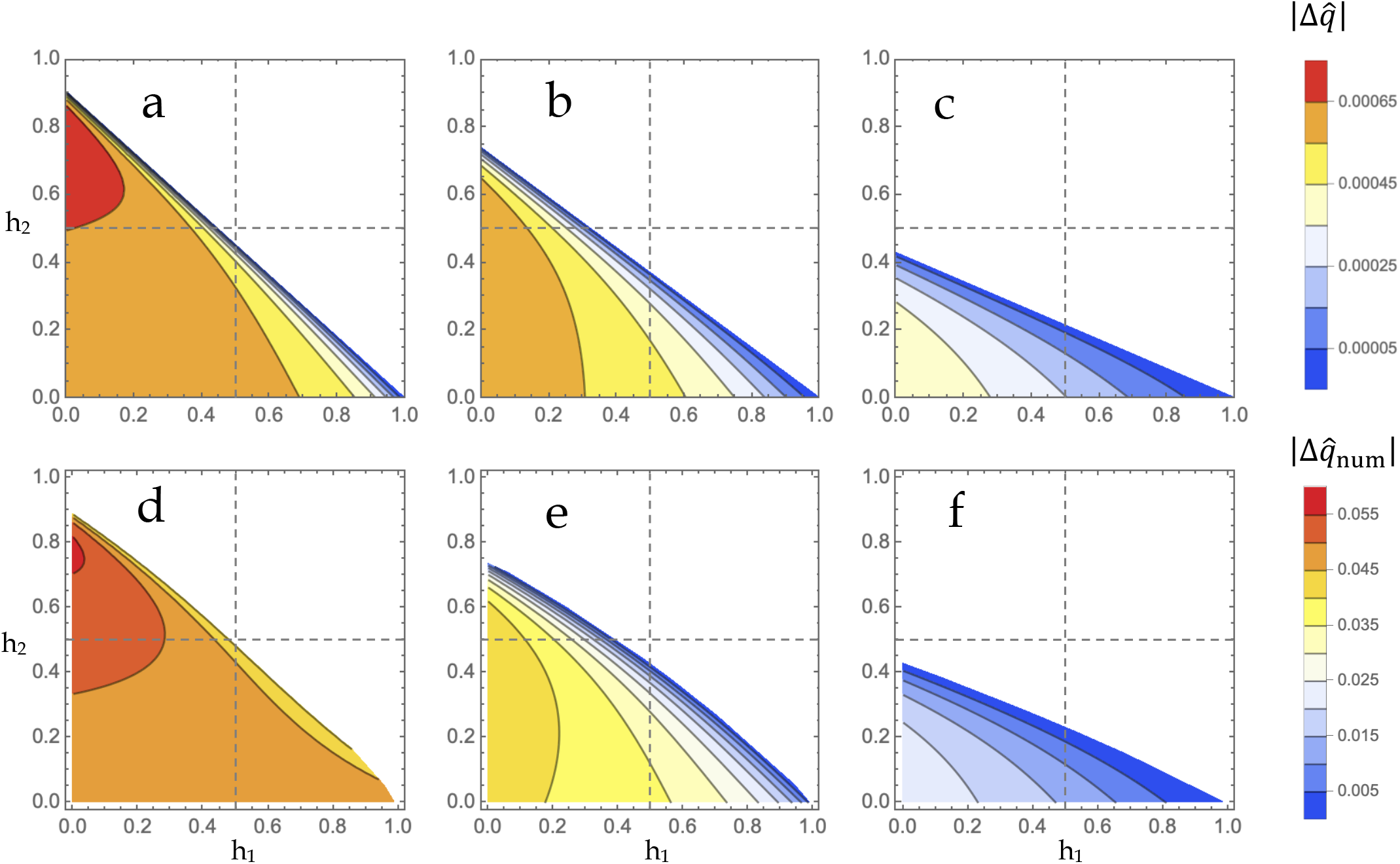
Amplitude of allelic oscillation in a bivoltine equilibrium. (a-f) Each point in the contour plots indicates the absolute value of the change in allele frequency between seasons 1 and 2 during the equilibrial cycle according to Eqn 20. (a-c) Weak selection plots, Eqn 20. (a) s_1_ = 0.525%, s_2_ = 0.475%, (b) s_1_ = 0.575%, s_2_ = 0.425% (c) s_1_ = 0.7%, s_2_ = 0.3%. (d-f) Strong selection plots, numerical. Allelic oscillation magnitudes were obtained for 2,101 parameter sets sampled from the region *P*. Recursions for the single-locus biallelic model were iterated for 10^4^ generations starting from an initial q = 0.5. (d) s_1_ = 32%, s_2_ = 35%, (e) s_1_ = 26%, s_2_ = 35% (f) s_1_ = 15%, s_2_ = 35%.

### Evolution of dominance

Upon establishment of the equilibrium cycle 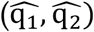, alleles outside of the set {A, a} may subsequently evolve by virtue of their effects on fitness. Consider a rare mutant (a_mut_) that is descended from the a-allele but which has a different season-specific dominance 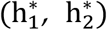 as compared to the resident population (h_1_, h_2_); selection coefficients remain unaltered.

The equations for the 3-allele system {A, a, a_mut_} are provided in Appendix 3.

To first-order in the selection coefficients, the fitness of a_mut_ upon introduction into the population is:

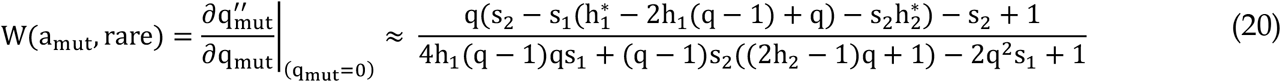

Assuming s_1_ ≤ s_2_, I consider three types of mutant effects and state the required conditions for allelic invasion into the resident state 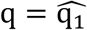:

**Case 1** Season-1-limited modification 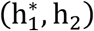:

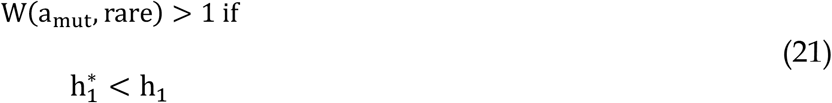

**Case 2** Season-2-limited modification 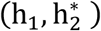:

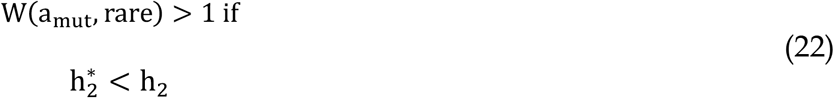

**Case 3** Season-specific modification 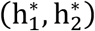:

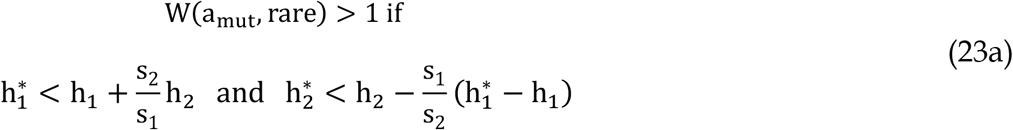

Or equivalently:

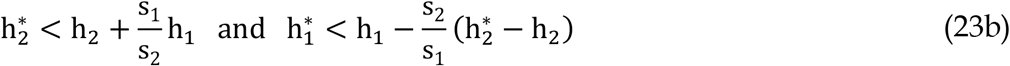

An increase in one season’s dominance coefficient requires a decrease in that of the other season. For these opposing changes, the conditions permit an increase to season 1 dominance that is larger than the magnitude of reduction in the season 2 coefficient; this is sensible given that selection is assumed to be less intense in season 1. In addition to oppositely directed changes, case 3 promotes seasonally-concordant reductions of dominance.

To the extent that mutants consist predominantly of seasonal-limited modifications (cases 1 and 2), the long-term evolutionary trend is toward the establishment of complete beneficial reversal 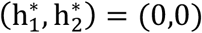, where the fitness of the heterozygote equals 1 in each season (Figure 5). Season-specific modification (case 3) allows for detours from this directional trend toward (0,0) and may initially cause a series of reversals between the two categories of constant dominance schemes, where the reversal in this case occurs over an evolutionary timescale rather than with the season (Figure 5). Nevertheless, the long-run trajectory in this case is also aimed at complete beneficial reversal, which, uniquely among resident phenotypes, is stable to the invasion of any possible mutants.

**Figure 5.**
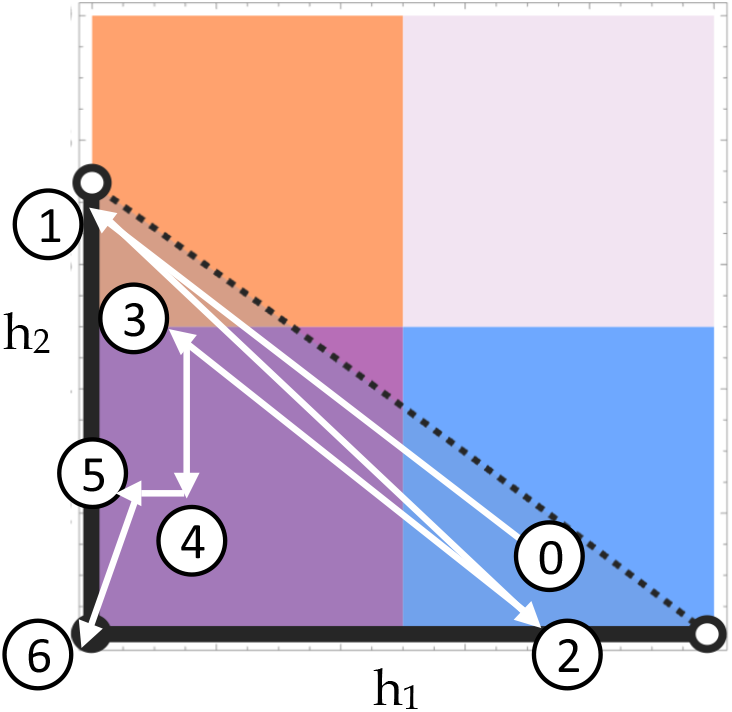
Hypothetical trajectory of dominance modification. Starting from a seasonally balanced equilibrium of the alleles {A, a} characterized by non-reversing dominance (0), successive replacements by mutant alleles with non-resident (h_1_,h_2_) schemes result in new temporary polymorphic equilibria (arrow ends). The full course of allele replacements may involve season-limited modifications (3→ 4 and 4→ 5) as well as season-specific changes (slanted arrows). Upon reaching the origin (6; complete beneficial reversal), the population is stable to any further modification.

[The same invasion conditions as above are obtained when considering the 3-allele system {A, A_mut_, a} for the dynamics of an A-descended mutant (Mathematica notebook file)].

The magnitude of selection on modifier mutants is typically sizeable. In Figure 6, I consider three examples of modification,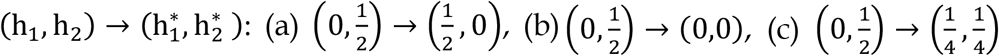. The resident population in (a)-(c) is assumed to exhibit complete A-allele dominance in season 1 and additivity in season 2. In scenario (a), the modifier alters the resident scheme by imposing additivity in season 1 and complete a-allele dominance in season 2. In scenario (b), the modifier directly establishes complete beneficial reversal, while in scenario (c), the modifier evolves an intermediate level of beneficial reversal. For the weak range of resident selection coefficients (s_1_,s_2_ ∼ 10^−4^ to 10^−3^), the effective selection coefficient favoring modifiers (*s*_*m*_, equal to the invasion fitness minus one) is greater than 10^−5^ for almost all of parameter space and can exceed 10^−3^ (Figure 6). Asymmetry of resident selection coefficients (s_1_, s_2_) favors modifier evolution with greater intensity than symmetric values; surprisingly, *s*_*m*_ can exceed the values of s_1_,s_2_ under strong resident asymmetry (Figure 6, red areas). The direct evolution of complete beneficial reversal (scenario b) is most favored among the three possibilities, while the seasonally non-reversing change (scenario a) generally has the second highest invasion fitness, and partial dominance reversal (scenario c) ranks third.

**Figure 6.**
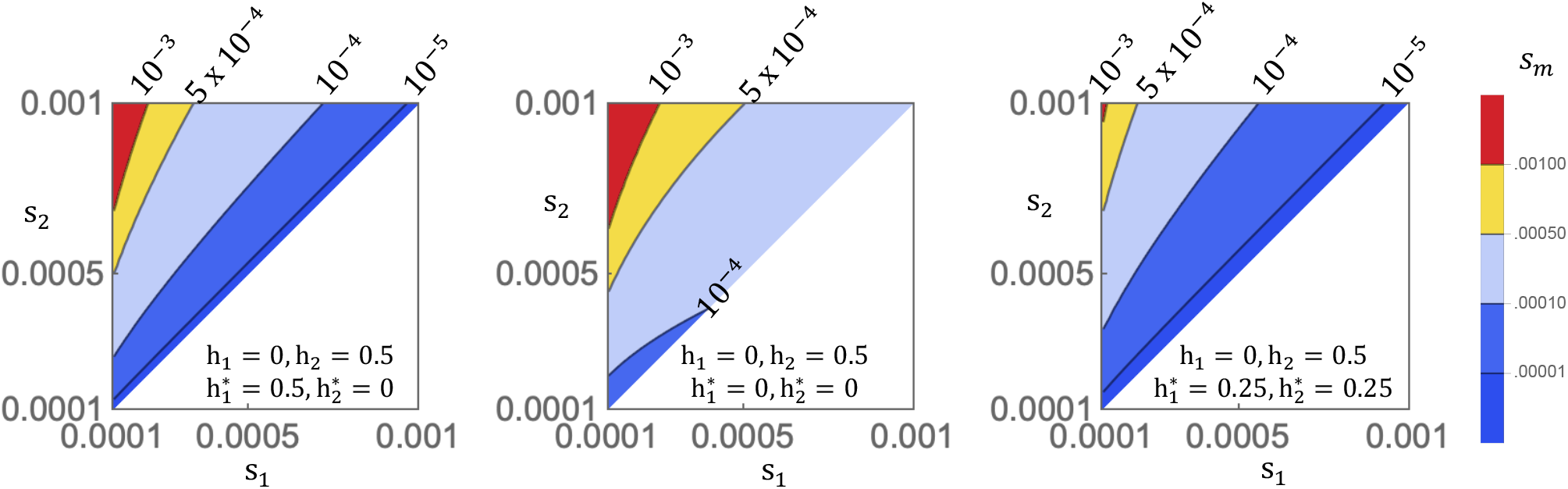
Strength of modifier selection. The value of *s*_*m*_, the effective selection coefficent favoring modifiers, is plotted for weak regimes of (s_1_, s_2_) in the resident population and assuming the indicated state of resident dominance (h_1_,h_2_) and the mutant effects of the modifier allele (h_1_*, h_2_*).

## Discussion

Having analysed the size of geometrically-overdominant parameter space, as well as its distribution over the major categories of season-specific dominance, I conclude the following: (1) seasonal fitness cycles are broadly favorable for the maintenance of polymorphism, and (2) much of the stabilizing potential owes to non-reversing allelic dominance. These conclusions apply for general intensities of selection and counter the notion that the assumptions required for temporally varying selection to act as a balancing mechanism are improbable. Moreover, these results challenge the role of dominance reversal as the primary stabilizing force acting on oscillating allelic variation. Random sampling of dominance parameters readily results in an annual heterozygote advantage (with 10-50% probability) and among these stabilizing schemes one-quarter to one-half involve constant directions of dominance. While the mutational process is unlikely to be uniform over the dominance unit square, mutation rates most likely favor non-reversing schemes over reversals and therefore such considerations would seem to further elevate the role of constant-directional dominance in maintaining polymorphism.

The bivoltine model of allele frequency change yielded additional insights into the characteristics of seasonally fluctuating equilibria. The model suggests that the evolution of large fluctuations is permitted broadly across most dominance schemes (Figure 5d-f), with the main limiting factor being the magnitude of seasonal fitness changes (s_1_, s_2_) and, secondarily, their symmetry: values near parity result in the largest possible swings. While oscillation is maximized by constant dominance, it remains the case that much of the polymorphic region (*P*) is associated with sizeable oscillation. Together with the invasion analysis of intralocus modifiers, the dynamical results suggest the following scenario for the history of seasonally balanced polymorphisms. Given an initially monomorphic locus, mutations with seasonally antagonistic effects (relative to the wild-type allele) originate in the population, and given realistic assumptions about mutation rates, the allele will tend to display partial dominance with a non-constant magnitude (i.e. one of the non-reversing quadrants of Figure 1). Over time, allele-specific expression changes may accumulate via divergence in the cis-regulatory regions of the two alleles, resulting in various possible changes including transient reversals in constant dominance (states 0, 1, and 2 in Figure 5) as well as changes that bring the dominance pattern closer to its ultimate (and optimal) fate: complete beneficial reversal. Such considerations provide the basis for predicting that the age of a seasonally balanced polymorphism is positively correlated with the proximity of its dominance scheme to maximal beneficial reversal.

In contrast to recent work (Wittmann et al. 2017, Bertram and Masel 2019, Wittmann et al. 2021, present article), the classical treatments of deterministic cyclical selection (Haldane and Jayakar 1963, Hoekstra 1975) avoided the decomposition of heterozygous fitness into dominance and selection parameters in their formulation of the model. Sufficient conditions giving rise to protected polymorphism (Levene 1953, Prout 1968) were found to be neatly expressible in terms of relative viabilities (Haldane and Jayaker 1963; Hoekstra 1975; Ewing 1977, 1979; Reinhold 1999, 2000), but ultimately such a formulation has limited informativeness with regard to questions about the genetic architecture of seasonal fitness and how this architecture is modified over time. After all, these matters depend partly on an understanding of how allelic dominance shapes seasonal fitness and so reformulating the geometric mean fitness criterion in terms of fundamental parameters is essential (Wittmann et al. 2017, Bertram and Masel 2019, Nagylaki 1975).

The “segregation lift” model of Wittmann et al. (2017) tackled such issues by focusing on two scenarios in a multilocus phenotypic context: (i) non-epistasis among loci and (ii) diminishing returns epistasis. They found that either scenario was permissive to the establishment of polymorphism with minimal or mild magnitudes of dominance reversal (with respect to a phenotypic score that is additive across loci). Wittmann et al. (2017) suggest a heuristic for determining whether seasonal varation can establish among a pair of alleles: namely that (h_1_+h_2_) < 1, i.e. that the sum of season-specific coefficients is less than one. As a matter of fact, this heuristic only works under perfect equality of the selection coefficients (s_1_=s_2_) and generally fails otherwise. Equations 4-5 are a superior replacement for general values of s_1_,s_2_ (with *m* = *n*; else use Equations 8 and 10 in asymmetric scenarios). An interesting argument was made by Bertram and Masel (2019) regarding the contrast between the stabilizing properties of beneficial reversal and the stabilizing properties of “cumulative overdominance” (Dempster 1955), also called incomplete dominance. For clarity, incomplete dominance here refers to schemes with constancy in both allelic direction and partial magnitude (i.e. the negatively sloped diagonal of the unit square).

Bertram and Masel’s (2019) argument emphasizes that beneficial reversal is broadly capable of establishing seasonal fluctuations across all selection intensities. Cumulative overdominance, on the other hand, is rather unsuited to the establishment of fluctuations at weak and moderate selection intensities, and therefore it establishes only large-amplitude oscillations. Whereas dominance reversal causes seasonal fitness to be composed of a mixture of weak, moderate and strong variation (presumably weighted toward weak alleles by the mutational process), cumulative overdominace would cause seasonal phenotypes to be predominantly constituted of large-effect alleles.

The results of the present article emphasize that the genetic basis of seasonal evolution is predicted to owe to a mixture of effect sizes and modes of gene action, and so no special relation/category of dominance, be it cumulative overdominance or beneficial reversal, is expected to constitute an exclusive role in stabilizing variation. Rather, seasonal balancing is predicted to involve (1) alleles that exhibit a range of selection coefficients and (2) heterozygous genotypes that are characterized by a distribution of season-specific dominance coefficients (Figure 2b, Box A1, Figure S1). I argue that cumulative overdominance fares poorly as a general explanation of sizeable seasonal fluctuations in large part because perfect correspondence to the complement relation h_1_ = 1-h_2_ is an implausible pattern of allelic dominance with respect to fitness. Let’s suppose that the season 1 coefficient takes some intermediate value of A-allele dominance, e.g. h_1_ = 0.22. On what basis do we form our expectation that the season 2 coefficient ought to be h_2_=0.78? After all, h_2_ refers to the allelic dominance in a rather different selective environment (e.g. winter vs. summer) and it is too special of a relation to expect that both direction and magnitude be preserved. The typical input by the mutational process is surely one in which partial magnitudes are in general uniquely valued. Allowance for this differentiation in seasonal magnitudes is precisely the emphasis of the present results, which show that much of the unit square contributes to polymorphism even apart from the beneficially reversing quadrant that was emphasized in Wittmann et al. (2017). To my knowledge, neither the distribution of season-specific dominance coefficients nor the contribution of small oscillations is known in any natural system; results thus far emphasize the observation of large seasonal changes in allele frequency across the genome (descriptive: Bergland et al. 2014, Machado et al. 2021; experimental: Rudman et al. 2022; see Johnson et al. 2023 for a recent review).

Situating the results in the broader topic of antagonistic pleiotropy (Hedrick 1999), my conclusions are sympathetic to earlier work that looked beyond the special dominance relations (e.g. additivity, incomplete dominance, symmetric dominance reversal). Curtsinger et al. (1994) sampled a rough grid of values from the dominance unit square in their numerical investigation of two-trait antagonistic pleiotropy. Despite finding a substantial proportion of stable parameter sets across the unit square, they dismissed as implausible that antagonistic pleiotropy promotes variation based on empirical results showing that the variance of dominance components in quantitative genetics experiments is typically small. Fry (2010) cautioned against concluding from narrow assumptions on autosomal dominance that sexual antagonism should be concentrated on the X chromosome; investigating numerical cases with partial dominance favoring the more fit allele in each sex, he concluded there exists a broader basis for the maintenance of genetic variation (see also Jordan and Charlesworth 2012). Van Dooren (2006) formulated an adaptive-dynamics model to investigate the likelihood of protected polymorphism in a multidimensional phenotypic context. He concluded that antagonistic pleiotropy and trait-specific dominance were necessary for polymorphism, and did not necessarily require beneficial reversal. Furthermore, he states that numerically the maximum possible ratio of reversing to non-reversing dominance (among stabilizing parameter sets) “appears to be 3” (p. 2001). The present study explains why this ratio appears as an analytical result in the two-season model (Figure 2b), and I likewise conclude that polymorphism owes to both categories of dominance. More recently, Spencer and Priest (2016) analyzed a modifier model of sexual antagonism, concluding that the sexes are expected to evolve dominance in opposing directions, accordant with similar conclusions arrived here in a seasonal context. The large amounts of both dominance variance and sex-by-dominance variance discovered in the seed beetle Callosobruchus maculatus (Grieshop and Arnqvist 2018), a model of sexual antagonism, testifies to the potential for widespread violations of constant-magnitude assumptions when dealing with antagonistic selection. Otto and Bourguet (1998) originally argued that balanced polymorphisms at intermediate frequencies were ripe for dominance modication. The intensity of selection on modifiers derived for the seasonal model (Figure 6) shores up this general conclusion regarding the unrestrictive prospects for the evolutionary adaptation of the genetic system to balanced polymorphism.

## Supporting information

proofs of areas

mathematica file

## Acknowledgments

I thank Rafael Guerrero, Paul Schmidt, Lucia Ramirez, Dmitri Petrov, and Alan Bergland for helpful discussion. I also thank Enrique Schwarzkopf and EvoGen seminar participants at NCSU. This work was supported by NIH/NIGMS grant no. R35GM147107 and NIH grant nos. R01GM100366 and R01GM137430.

## Appendix 1

The quantitative contribution of each quadrant to the size of *P* is calculated by: (1) solving for the absolute areas of the quadrant-specific subregions of *P* (Supplemental Section 1), and then (2) dividing each value by Area(*P*) (Equation 6) (Box A1). Cases i-iii are distinguished by the number of quadrants with positive areas.

### Box A1.

**The relative contributions of the four major categories of season-specific dominance to the potential for biallelic seasonally balanced polymorphism (*m* = *n*).**

The area of a subregion relative to the size of *P* is given by RelArea 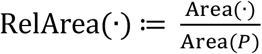

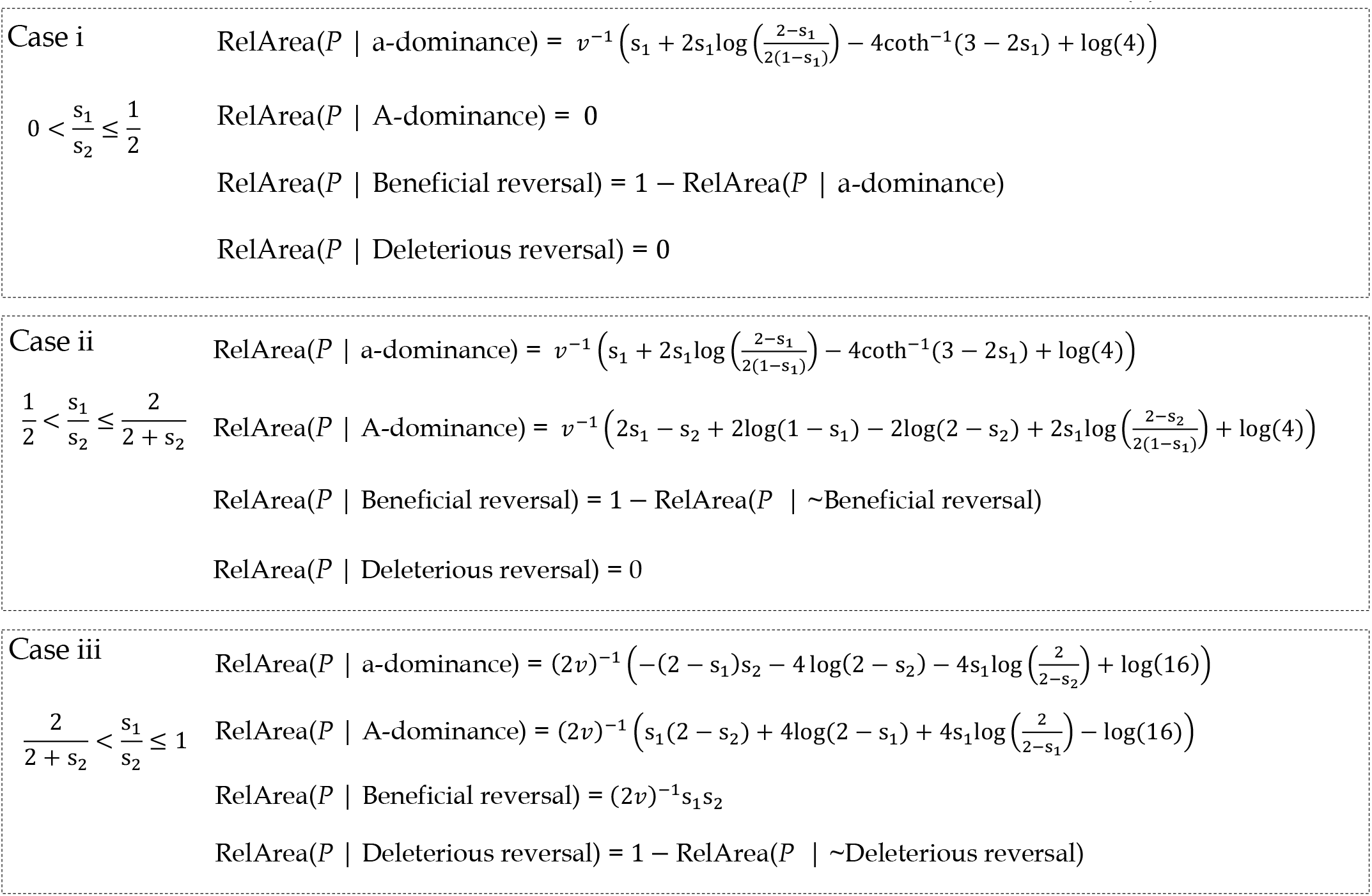

where *v* = 2(s_1_ + (1 – s_1_)log(1 – s_1_)). See Supplemental Section 1 for proofs.

## Appendix 2

The classical bivoltine model (Haldane and Jayakar 1963) can be written in terms of the fitnesses in Table 1, resulting in the system of equations for season 1:

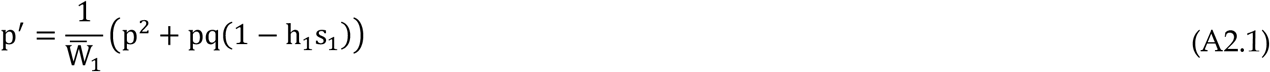

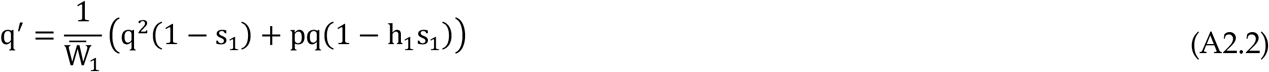

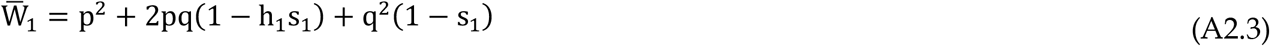

and season 2:

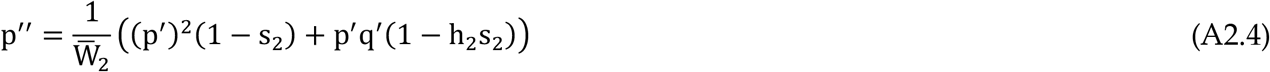

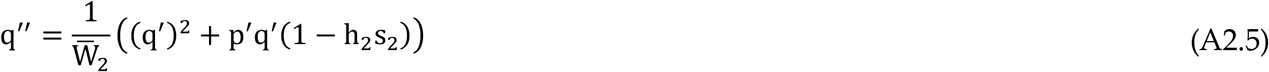

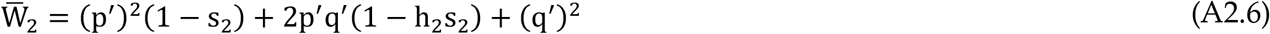

where p, q is the frequency of A, a, respectively, at the start of season 1 and p + q = 1. The ratio of allele frequencies is q/p for the initial generation, and is equal to q ^″^ /p ^″^ following an annual cycle. The ratio of these ratios is equal to the relative annual fitness of the a-allele (w_a_):

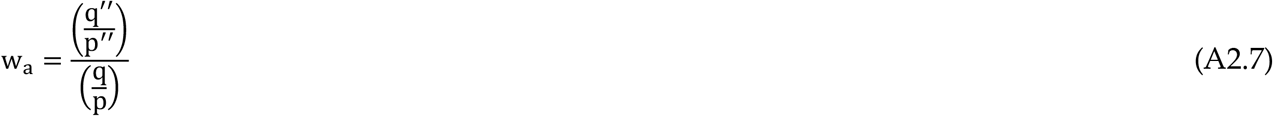

## Appendix 3

Evolutionary changes in the 3-allele system {A, a, a_mut_} are governed by the following equations, which employ mutant dominance relations 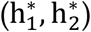 for Aa_mut_ heterozygotes.

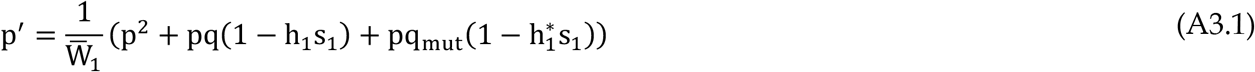

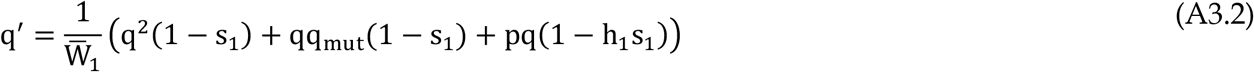

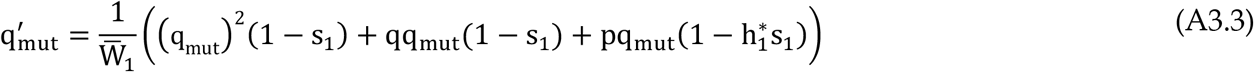

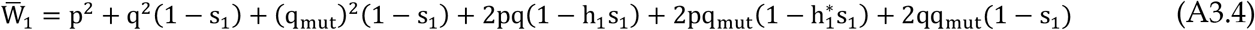

and for season 2:

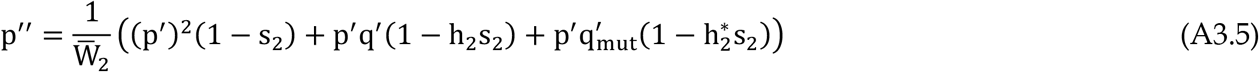

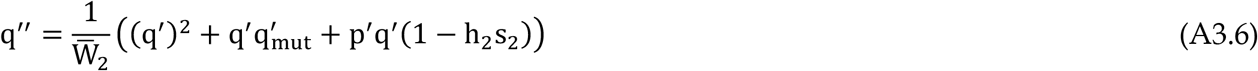

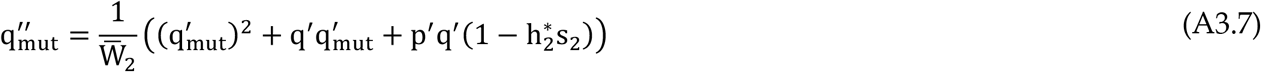

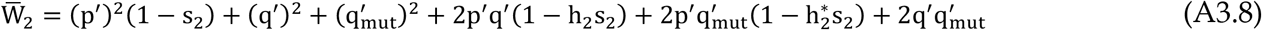

where p + q + q_mut_ = 1.

