## Supplementary material for "Season-specific dominance broadly stabilizes polymorphism under symmetric and asymmetric multivoltinism": proofs of areas

by Evgeny Brud

The absolute areas of the quadrant-specific subregions of $P$ are derived by cases. These areas are each divided by the total area of $P$ in order to obtain the relative areas of Box I.

Case i: $0 < \frac{s_1}{s_2} \leq \frac{1}{2}$

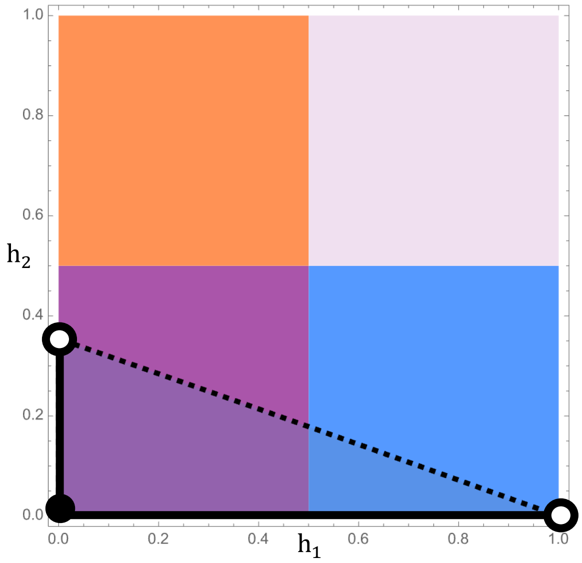


$$\text{Area(P)} = \int_0^1 h_{2,max}\, dh_1$$

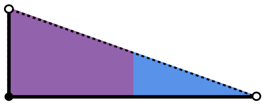

$$= \frac{(1 - s_1)\log(1 - s_1) + s_1}{s_1 s_2}$$

$$\text{Area(P | a- dominance, case 1)} = \int_{1/2}^{1} h_{2,max}\, dh_1$$

$$= \frac{\left(s_1 + 2s_1 \log\left(\frac{2 - s_1}{2(1 - s_1)}\right) - 4\coth^{-1}(3 - 2s_1) + \log(4)\right)}{2 s_1 s_2}$$

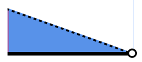

Area(P | beneficial reversal, case 1)

$$= \text{Area(P)} - \text{Area(P | a} - \text{ dominance, case 1)}$$

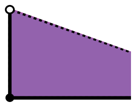

Case ii: $\frac{1}{2} < \frac{s_1}{s_2} \leq \frac{2}{2+s_2}$

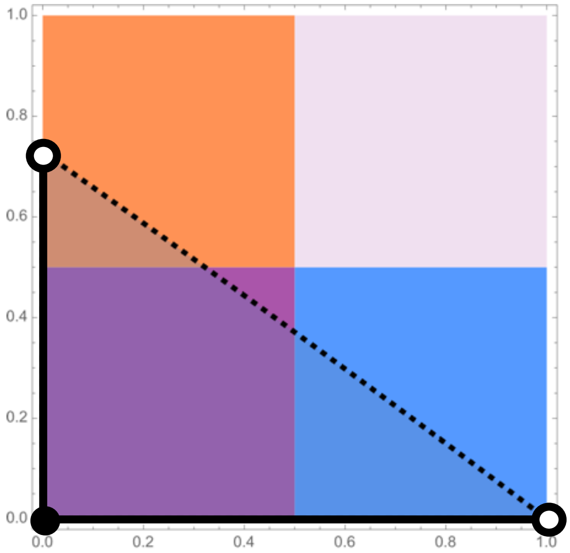


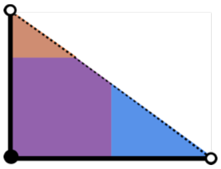

$$\text{Area(P)} = \int_0^1 h_{2,max}\, dh_1$$

$$= \frac{(1 - s_1)\log(1 - s_1) + s_1}{s_1 s_2}$$

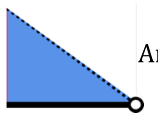

$$\text{Area(P | a- dominance, case 2)} = \int_{\frac{1}{2}}^{1} h_{2,max}\, dh_1$$

$$= \frac{s_1 + 2s_1 \log\left(\frac{2 - s_1}{2(1 - s_1)}\right) - 4\coth^{-1}(3 - 2s_1) + \log(4)}{2s_1 s_2}$$

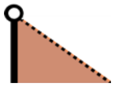

$$\text{Area(P | A- dominance,case 2)} = \int_{\frac{1}{2}}^{s_1/s_2} h_{1,max}\, dh_2$$

$$= \frac{\left(2s_1 - s_2 + 2\log(1 - s_1) - 2\log(2 - s_2) + 2s_1 \log\left(\frac{2 - s_2}{2(1 - s_1)}\right) + \log(4)\right)}{2s_1 s_2}$$

Area(P | beneficial reversal, case 2) =

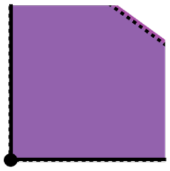 = 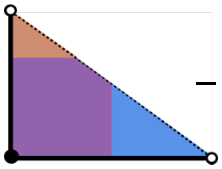 − 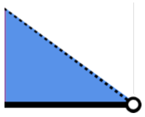 − 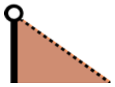

Case iii: $\frac{2}{2+s_2} < \frac{s_1}{s_2} \leq 1$

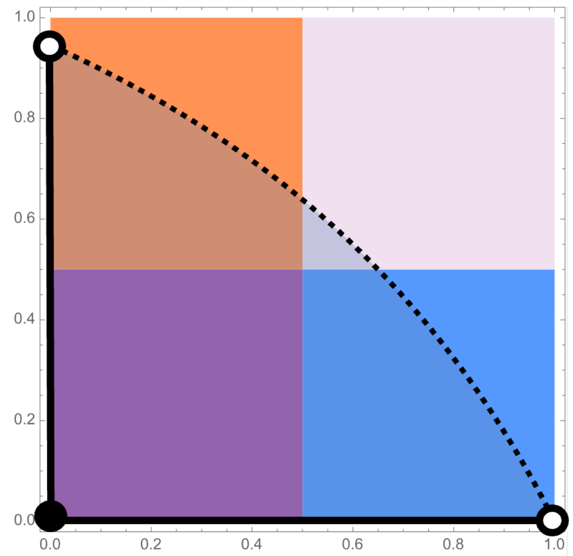


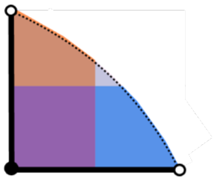

$$\text{Area}(P) = \int_0^1 h_{2,max}\, dh_1$$

$$= \frac{(1-s_2)\log(1-s_2)+s_2}{s_1 s_2}$$

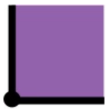

$$\text{Area}(P \mid \text{beneficial reversal, case 3}) = \frac{1}{4}$$

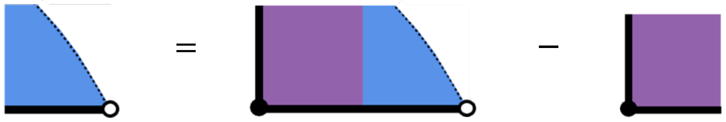

$$\text{Area}(P \mid a-\ \text{dominance, case 3}) \;= \int_0^{1/2} h_{1,max}\, dh_2 - \frac{1}{4}$$

$$= \frac{s_2 + 2\log(2-s_2) + 2s_1\log\left(\frac{2}{2-s_2}\right) - \log(4)}{2s_1 s_2} - \frac{1}{4}$$

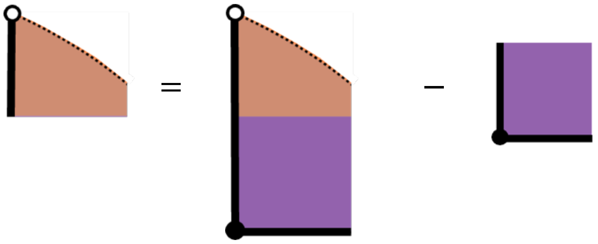

$$\text{Area}(P \mid A-\ \text{dominance, case 3}) \;= \int_0^{1/2} h_{2,max}\, dh_1 - \frac{1}{4}$$

$$= \frac{s_1 + 2\log(2-s_1) + 2s_1\log\left(\frac{2}{2-s_1}\right) - \log(4)}{2s_1 s_2} - \frac{1}{4}$$

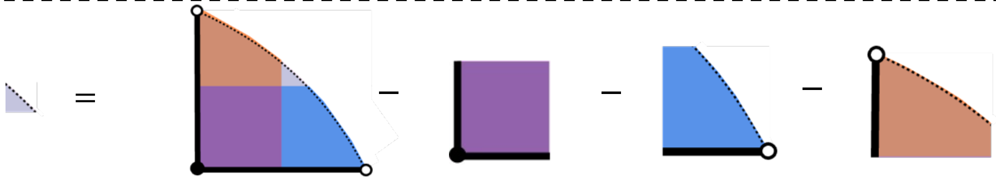

$= \text{Area}(P \mid \text{deleterious reversal, case 3})$
