## additional figures for "Season-specific dominance broadly stabilizes polymorphism under symmetric and asymmetric multivoltinism"

by Evgeny Brud

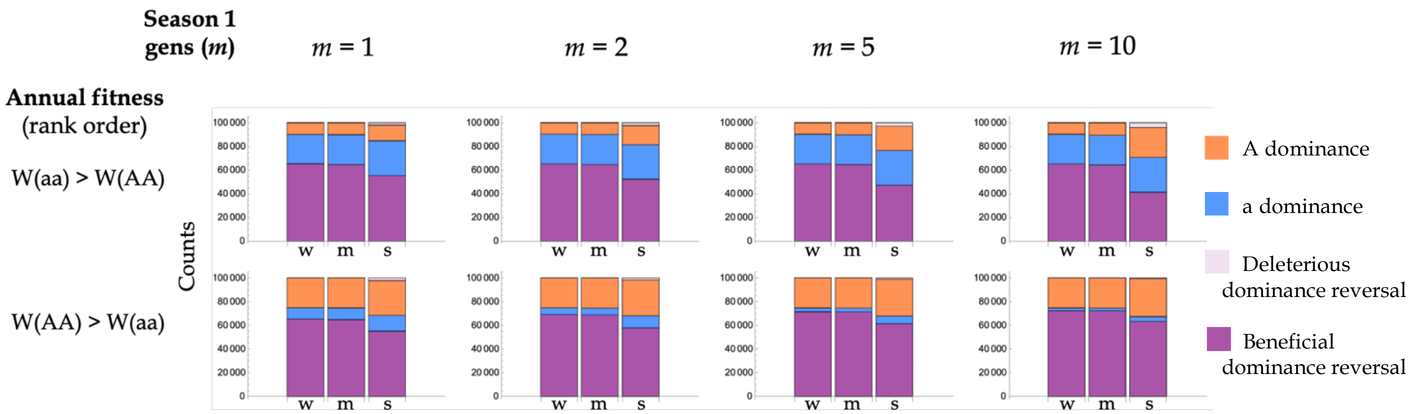


**Figure S1 Numerical distribution of stabilizing dominance schemes under symmetric and asymmetric multivoltinism.** Stacked bars indicate the counts for each category of season-specific dominance under weak (0-0.1%; w), moderate (0.1-10%; m), and strong (10-100%; s) ranges for the selection coefficients, assuming the region in Eqn 2 for the generation numbers $(m, n) = (m, 1)$ and random uniform sampling of $10^5$ points per parameter set. (Top row) Sampling assumes condition 7a for the rank order of homozygous annual fitnesses, i.e. W(aa)>W(AA). (Bottom row) Sampling assumes condition 7b, i.e. W(AA)>W(aa).

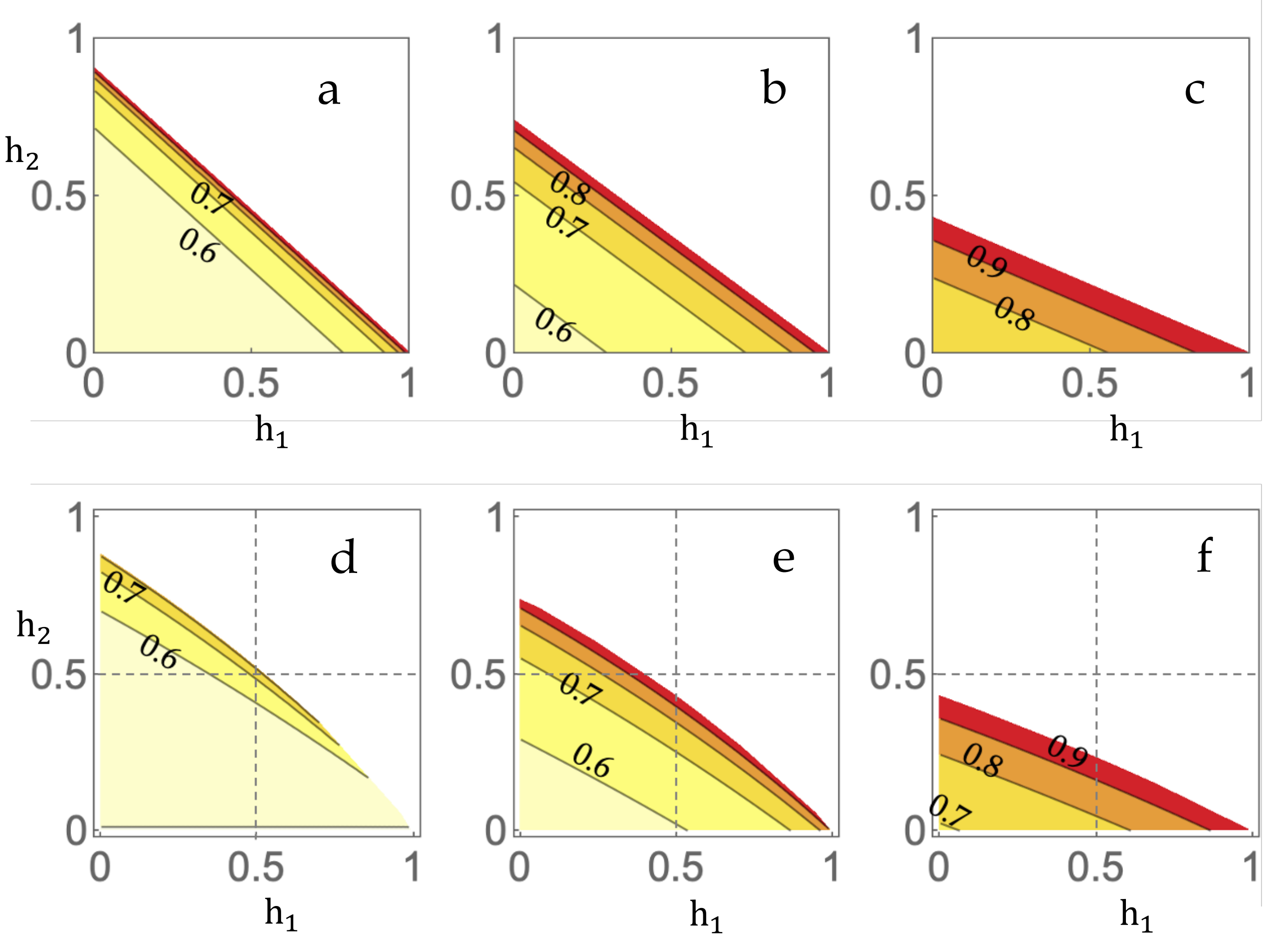


**Figure S2 Equilibrium a-allele frequency at the start of season 1 in a bivoltine population.** (a-c) The value of $\hat{q}_1$ is plotted for weak regimes of ($s_1$, $s_2$). (a) $s_1$ = 0.525%, $s_2$ = 0.475%, (b) $s_1$ = 0.575%, $s_2$ = 0.425% (c) $s_1$ = 0.7%, $s_2$ = 0.3%.(d-f) The numerical value of the a-allele frequency at generation $10^4$ ($\hat{q}_{1_{\mathrm{num}}}$) is plotted under strong selection regimes. The biallelic model was iterated from q = 0.5 at gen 0. (d) $s_1$ = 32%, $s_2$ = 35%, (e) $s_1$ = 26%, $s_2$ = 35% (f) $s_1$ = 15%, $s_2$ = 35%.
